# Real-time Metagenomic Analysis of Undiagnosed Fever Cases Unveils a Yellow Fever Outbreak in Edo State, Nigeria

**DOI:** 10.1101/572354

**Authors:** Fehintola V. Ajogbasile, Judith U. Oguzie, Paul E. Oluniyi, Philomena Eromon, Jessica Uwanibe, Samar B. Mehta, Katherine J. Siddle, Ikponmwosa Odia, Sarah M. Winnicki, Nosa Akpede, George Akpede, Sylvanus Okogbenin, Bronwyn L. MacInnis, Onikepe A. Folarin, Stephen F. Schaffner, Ephraim Ogbaini-Emovon, Oyewale Tomori, Chikwe Ihekweazu, Pardis C. Sabeti, Christian T. Happi

**Affiliations:** African Center of Excellence for Genomics of Infectious Diseases (ACEGID), Redeemer’s University, Ede, Osun State, Nigeria.; Department of Biological Sciences, College of Natural Sciences, Redeemer’s University, Ede, Osun State, Nigeria.; Broad Institute of MIT and Harvard, Cambridge, Massachusetts, USA.; Beth Israel Deaconess Medical Center, Division of Infectious Diseases, Boston, Massachusetts, USA.; Center for Systems Biology, Department of Organismic and Evolutionary Biology, Harvard University, Cambridge, Massachusetts, USA.; Institute of Lassa Fever Research and Control, Irrua Specialist Teaching Hospital, Irrua, Edo State, Nigeria.; Department of Immunology and Infectious Diseases, Harvard T.H. Chan School of Public Health, Harvard University, Boston, Massachusetts, USA.; Nigerian Centre for Disease Control, Abuja, Nigeria.; Howard Hughes Medical Institute, Chevy Chase, Maryland, USA.

## Abstract

We report here the use of metagenomic sequencing to identify and characterize yellow fever virus (YFV) in a cluster of undiagnosed febrile patients from Edo State, Nigeria. Yellow fever, an acute mosquito-borne viral haemorrhagic fever, re-emerged in Nigeria in 2017 after decades of relative control.^1^ Once a major public health concern in the region,^2^ the prolonged absence of yellow fever has left a gap in knowledge about circulating virus and lowered clinical suspicion for a disease whose spectrum overlaps with endemic infections (*e.g.*, malaria and Lassa fever). In this context, metagenomic sequencing can be a powerful tool for identifying and tracking pathogens causing infectious disease.^3^

Plasma samples from thirteen epidemiologically-linked patients with unexplained, often fatal, fever were sent from Irrua Specialist Teaching Hospital to the African Center of Excellence for Genomics of Infectious Disease at Redeemer’s University for investigation. Unbiased metagenomic sequencing revealed YFV in seven samples, from which four genomes were assembled (two full and two partial). We confirmed this by RT-qPCR, which also detected YFV in two additional samples (Supplementary Table 1). No other viral pathogens were detected in any of the samples, including those negative for YFV (Supplementary Figure 2). YFV has a short period of viraemia,^4^ which may explain the absence of detectable virus in the four other patients.

To determine the source of the outbreak in Edo State, we compared these genomes to all previous African YFV sequences available in GenBank, using the portion of the genome (prM/E gene) most widely sequenced in historical samples. In a maximum likelihood phylogeny, the 2018 YFV sequences formed a tightly clustered clade (Figure 1), more closely related to samples from other West African countries (mean pairwise identity = 99·4%) than to earlier (1946–1991) Nigerian sequences (mean pairwise identity = 89·8%). This provides the first evidence that YFV responsible for the 2018 outbreak in Edo State does not descend directly from the Nigerian YFV outbreaks of the last century but is instead part of the broader diversity of current YFV in West Africa.

**Figure 1:**
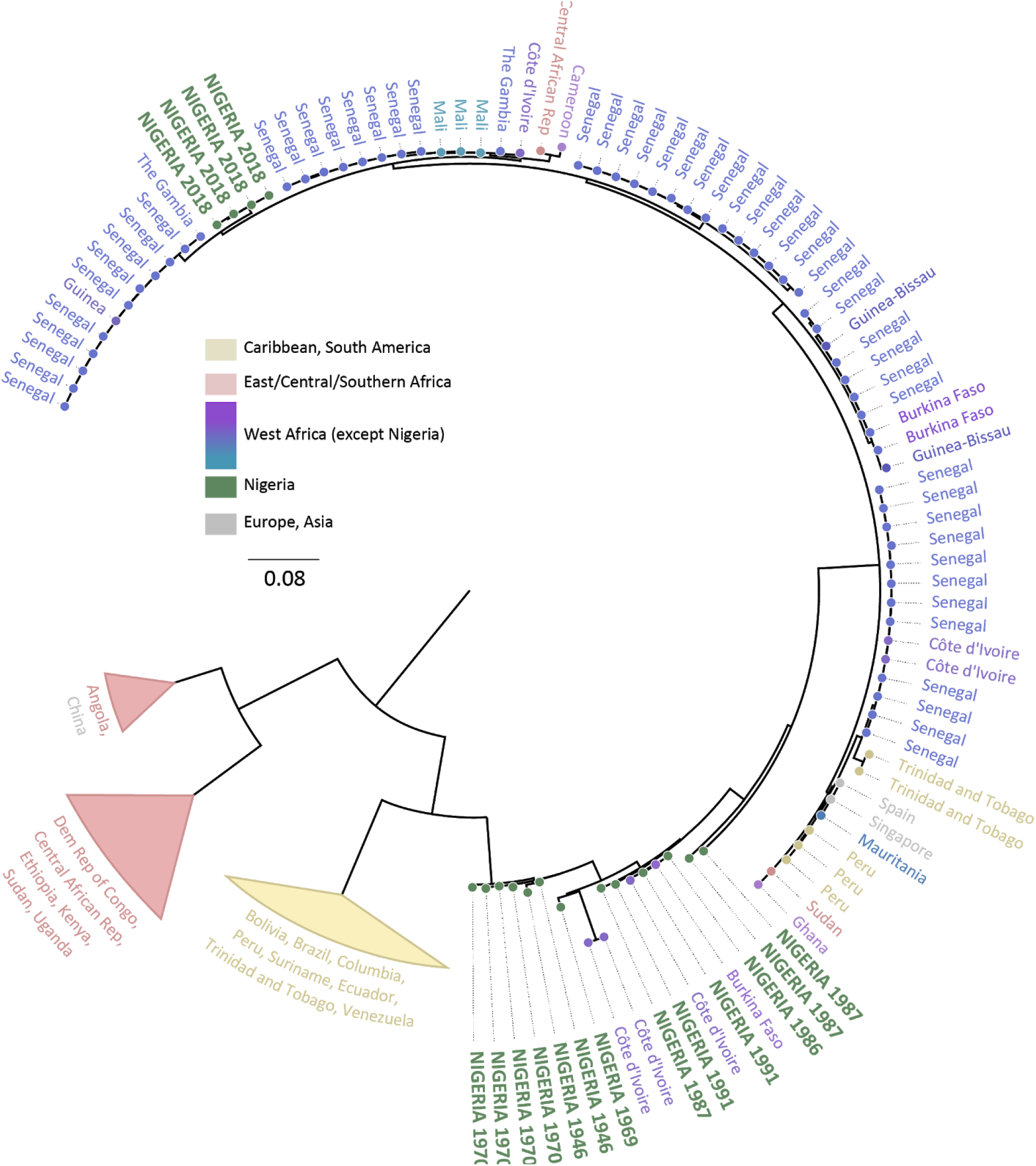
Maximum likelihood tree for a region of the prM/E genes from 205 YFV sequences.

We established the presence of YFV in Edo State within four days of receiving samples and shared this information immediately with the referring hospital and national health authorities. Based in part on these findings, the Nigeria Centre for Disease Control declared an outbreak in Edo the following day.^5^ Notably, these are the only sequence data reported from recent Nigerian YFV cases, and the first complete Nigerian YFV genomes from patient samples collected after 1950. This ability to rapidly identify and characterize a re-emerging virus – in an unusual cluster identified by local health officials – highlights the value of in-country genomics capacity. The integration of this capacity into the established, but siloed, pathogen-specific diagnostic platforms developed over the past 20 years provides exciting opportunities for public health surveillance.

## Acknowledgements

We appreciate the continuous support of ACEGID staff and the management of Redeemer’s University, and we thank our colleagues the Broad Institute, the Nigerian Center for Disease Control (NCDC), and the Irrua Specialist Teaching Hospital (ISTH). We specifically recognize the ongoing, invaluable dedication of the laboratory staff of the Institute for Lassa Fever Research and Control laboratory staff at ISTH: D. I. Adomeh, C. Aire, J. Agbukor, M. Airende, J. Aiyepada, P. Akhilomen, P. Ebhodaghe, B. Ebo, A. Ekanem, S. Ekhikhametalor, R. Enigbe, R. Esumeh, R. Giwa, G. Ignegbale, E. Muoebonam, G. Odigie, O. Omoregie, T. Olokor, G. Okonofua, E. Omomoh, R. Omiunu, B. Osiemi, J. Oyakhilome, and E.O. Yerumoh. We dedicate this work to the people in the affected communities of Edo State.

## Funding statement

This work is supported by grants from the National Institute of Allergy and Infectious Diseases, NIH-H3Africa (U01HG007480 and U54HG007480 to Redeemer’s University [Dr. Happi]) and a grant from the World Bank (project ACE019 to Redeemer’s University [Dr. Happi]). Dr Mehta is supported by the Division of Infectious Diseases at Beth Israel Deaconess Medical Center. Dr. Siddle is supported by a fellowship from the Human Frontier Science Program (LT000553/2016). Dr. Sabeti is an investigator supported by the Howard Hughes Medical Institute (HHMI).

## SUPPLEMENTAL TEXT

### Sample collection and testing

Thirteen patient samples were identified retrospectively after ISTH clinicians recognized a pattern in several patients with overlapping presentation, poor outcomes, and lack of clear diagnosis. The patients came from contiguous local government areas; their samples were initially referred to ISTH for Lassa fever molecular testing. All patient samples tested negative for Lassa virus by RT-qPCR. Eight patients had documented negative testing for malaria (either RDT or microscopy).

### Sample Preparation and sequencing

Plasma was inactivated in AVL and viral RNA extracted according to the QiAmp viral RNA mini kit (Qiagen) according to the manufacturer’s instruction. Extracted RNA was treated with Turbo DNase to remove contaminating DNA, followed by cDNA synthesis with random hexamers. Sequencing libraries were prepared using the Nextera XT kit (Illumina) as previously described^1^ and sequenced on the Illumina Miseq platform with a read length of 101 base pair paired end.

### SYBR based YFV qRT-PCR

Sybr green RT-qPCR was performed on a Roche LightCycler 96. Briefly, 3µl of RNA was used per reaction as a template for amplification. The sample was added to 7µl of reaction mixture containing 1·32µl of H_2_O, 5µl of Power Sybr master mix, 0·08µl of 125X reaction mix and 0·3µl sense and anti-sense primers. Real-time RT-qPCR amplification was carried out for 45 cycles of 48°C for 30 min, 95°C for 10 min, 95°C for 15 sec, and 60°C for 30 sec. Temperature ranges for the melt curves was 95°C for 15 sec, 55°C for 15 sec and 95°C for 15 sec. The yellow fever primer sequences have been published.^2^

### Metagenomic analysis of viral infections

We used Kraken^3^ to perform an initial taxonomic classification of all viral taxa present in the sample using a database that encompassed the known diversity of all viruses that infect humans, as previously described.^4^ To confirm and obtain de-duplicated counts of classified reads we then performed alignment to reference genomes of all viruses identified by Kraken as present in one or more samples. We performed alignment with Novoalign (http://www.novocraft.com) using the following stringent parameters: “-k -l 40 -g 40 -x 20 -t 100-r Random” and used Picard (http://broadinstitute.github.io/picard) to mark and remove duplicates. We did not find strong evidence of other viral infections beside YFV in these samples.

### Genome Assembly and Maximum Likelihood Phylogenetic Analysis

Viral genomes were assembled using viral-ngs v1.21.2^5, 6^ on DNAnexus platform, and MAFFT v7.310^7^ was used to align them with all African YFV genomes available in GenBank as of 19th January 2019 (including a small number of non-African sequences as an outgroup). Using Geneious 2019.0.4,^8^ an approximately 680bp of the prM/E region of the genome which was the most covered by existing GenBank sequences from Africa was extracted and used to infer a maximum likelihood tree using IQTREE v1.5.5.^9^ We used a Tamura-Nei nucleotide-substitution model with a gamma distribution of rate variation among sites^10^ and ultrafast bootstrapping.^11^

### Data Availability

The four sequences from this study are available on GenBank with Accession numbers: MK457700, MK457701, MK457702, MK457703.

### Bayesian Phylogenetic Analysis

Time-scaled Bayesian phylogenetic analysis was carried out using the Markov chain Monte Carlo (MCMC) algorithm implemented in the BEAST v1.10.4^12^ package with BEAGLE^13^ to improve run-time. The evolutionary and demographic process were estimated from the sampling dates of the prM/E sequences using a model that incorporated a General Time Reversible (GTR) + Gamma distribution (four categories) model with “(1+2),3” codon partitioning, an uncorrelated relaxed clock with log-normal distribution,^14^ and a Bayesian skyline coalescent tree prior distribution.^15^ All the Bayesian analyses were run for 200 million MCMC steps, with parameters and trees sampled every 10000 generations. Uncertainty of parameter estimates were assessed by calculating the Effective Sample Size (ESS) and the 95% Highest Probability Density (HPD) values, respectively using TRACER v1.6.0^16^ program. Maximum clade credibility trees summarizing all Markov chain Monte Carlo samples were generated with the use of TreeAnnotator v1.10.4^17^ software, with a burn-in rate of 10% and FigTree v1.4.4^18^ was used to view and annotate the MCC tree. This analysis used the same 680bp of the prM/E region of the YFV genome as was used for the maximum likelihood approach above, possibly adding a bias our tMRCA estimates.

---

**Supplementary Figure 1:**
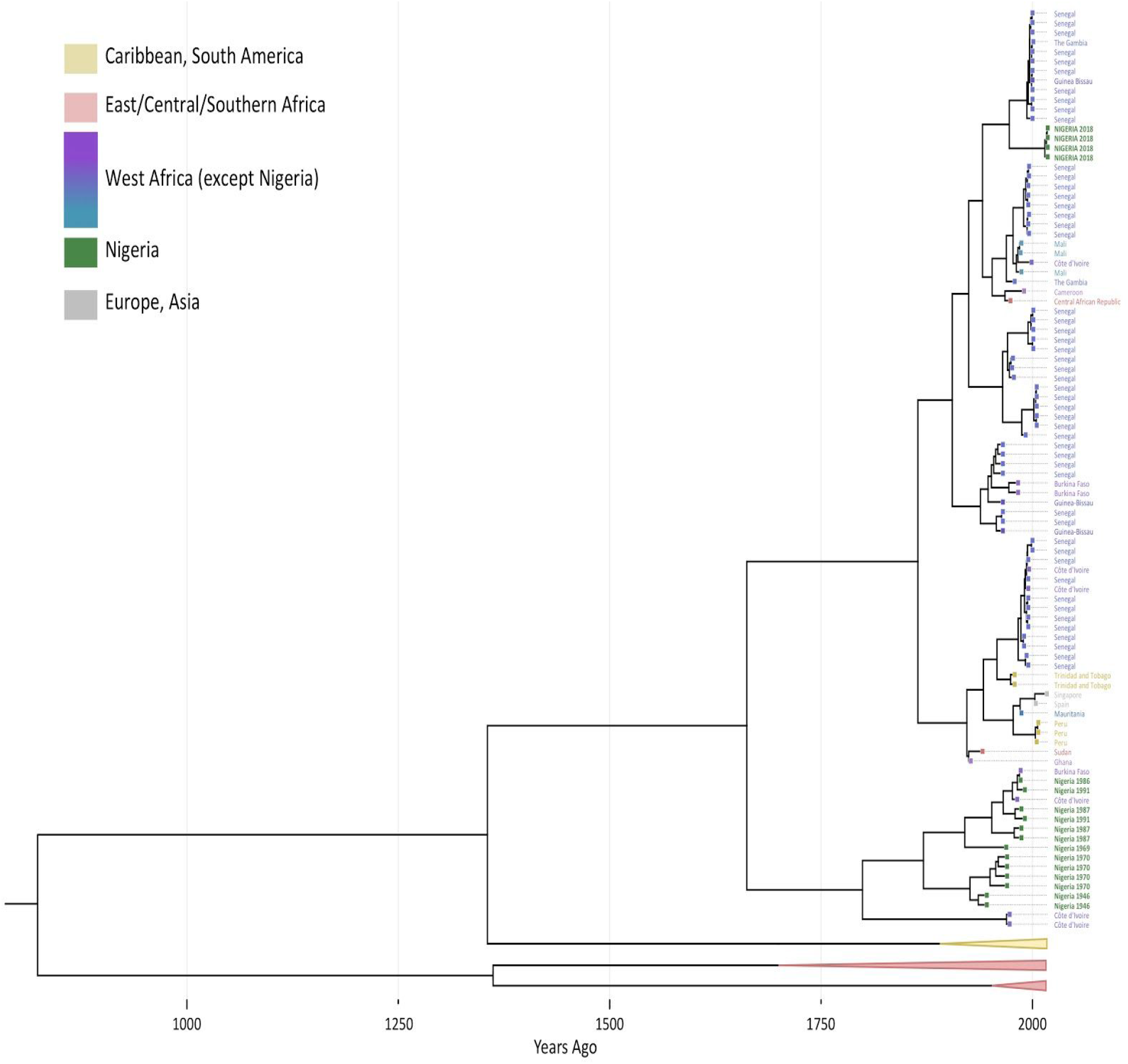
Time-scaled Bayesian MCC phylogenetic tree for a region of the prM/E gene of 205 YFV sequences. Sequences were aligned using MAFFT. The tMRCA of the four (4) 2018 sequences from Nigeria is 47 years, 95% HPD interval [26,74], while the tMRCA of the pre-1991 sequences from Nigeria is 127 years, 95% HPD interval [88, 175].

**Supplementary Figure 2:**
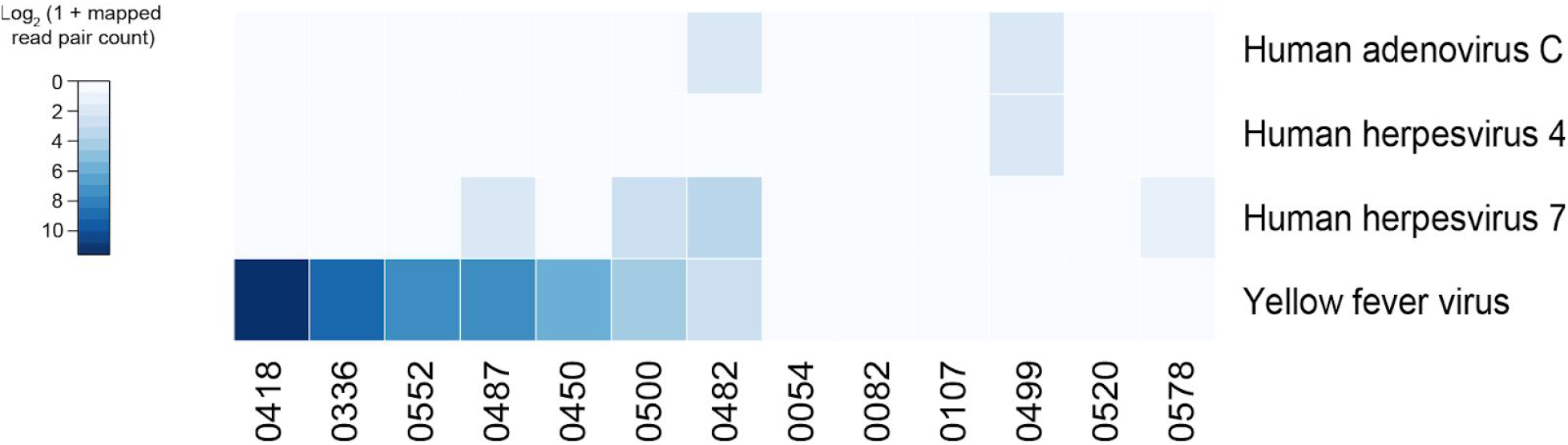
Virus detection by metagenomic sequencing. Heatmap shows viral species where at least one sample produced reads that aligned to the viral RefSeq genome. Detected species were first filtered using Kraken. Yellow fever virus was the only virus found to be present in any sample at appreciable levels.

**Supplementary Table 1:**
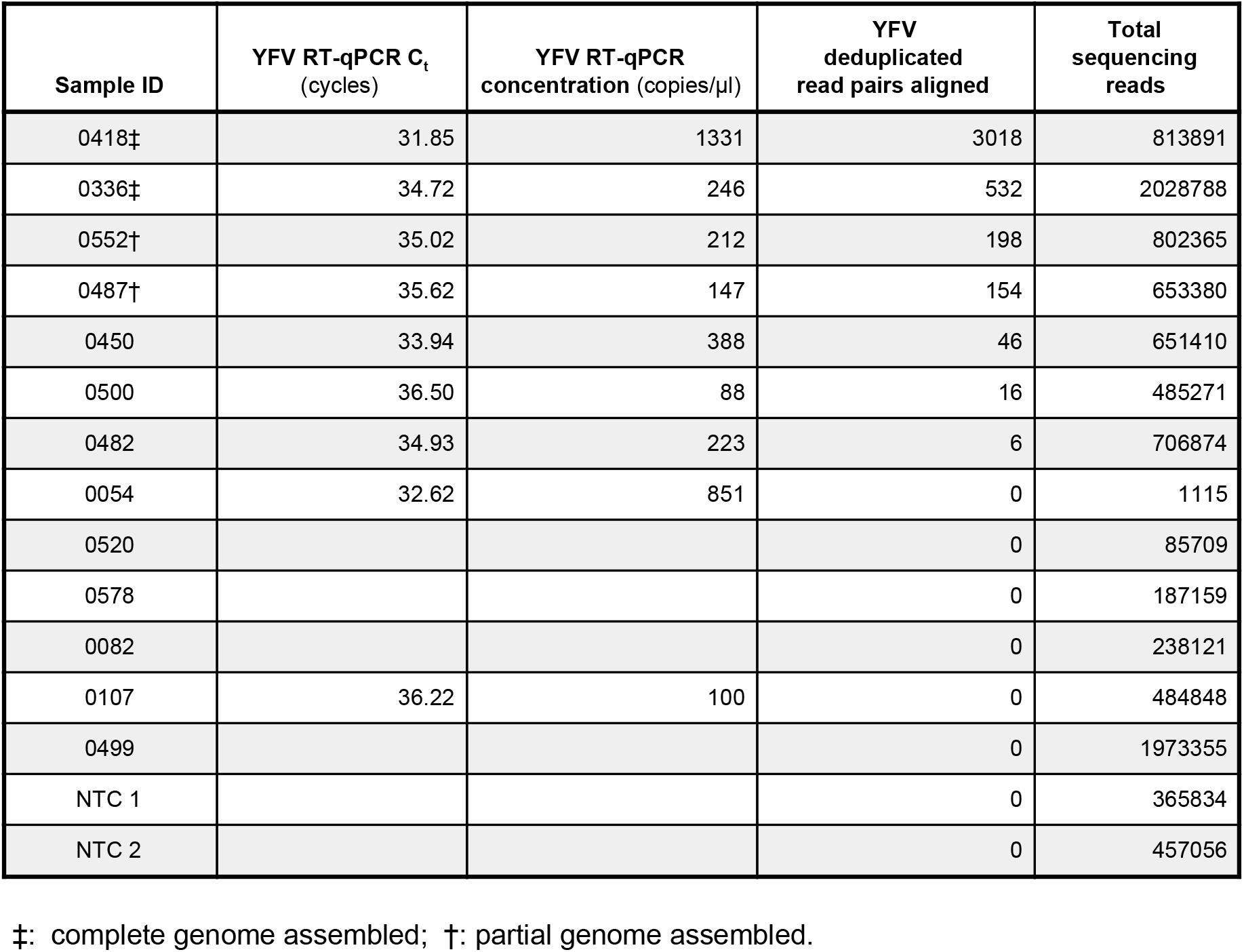
Yellow fever RT-qPCR & sequencing results.

